# Effects of Malaysian strains of *Toxoplasma gondii* on behaviours and their possible risk in schizophrenia-like rat model

**DOI:** 10.1101/2020.12.10.418301

**Authors:** Mohammed Nasiru Wana, Malaika Watanabe, Samaila Musa Chiroma, Ngah Zasmy Unyah, Sharif Alhassan Abdullahi, Onesimus Mahdi, Ashraf Ahmad Isa Alapid, Shariza Nordin, Rusliza Basir, Mohamad Aris Mohd Moklas, Roslaini Abd. Majid

## Abstract

*Toxoplasma gondii* (*T. gondii*) is a protozoan parasite that reside majorly in the brain of its intermediate host. *T. gondii* infected rodent’s shows some degree of behaviour deficits, while *T. gondii* infection in humans is associated with psychiatric problems such as schizophrenia. The present study aimed to evaluate the effects of Malaysian strains of *T. gondii* on rats. Forty five, four weeks old, male Wistar rats were used. The rats were assigned into five groups: two control groups (CG1 and CG2) and three experimental groups (EG1, EG2, EG3). CG1 rats received phosphate buffered saline (PBS), CG2 received MK-801 (as a model for schizophrenia), EG1, EG2, EG3 received orally 5 × 10^3^ single *T. gondii* oocysts strain of type I, type II and type III respectively. After infection, all the five groups of rats were tested for *T. gondii* antibodies at two weeks post-infection (PI). Behavioural tests of exploratory activity (open field) and spatial learning and memory retention (Morris water maze) were performed on the ninth and tenth weeks PI followed by histological staining of rat brain. *T. gondii* IgM antibodies were detected in EG1, EG2 and EG3, but not in CG1 and CG2. The behaviour results demonstrated that rats from CG2, EG1, EG2 and EG3 had increased in their locomotor activities and memory deficits compared to control, while learning remain intact. Moreover, tissue cysts were found widely distributed exclusively in the whole brain of EG1, EG2 and EG3 without tropism. These findings taken together, implies that Malaysian strains of *T. gondii* are implicated in some causes of behaviour changes that are responsible for schizophrenia-like conditions if humans were infected.

## 1. Introduction

*Toxoplasma gondii* (*T. gondii*) is a ubiquitous protozoan parasite which is acquired through ingestion of oocysts contaminated in food, water and vegetables (Dubey and Dubey 2010). Other routes include consumption of undercooked meat containing cysts or through placenta transmission from mother to unborn child (Afonso et al. 2013). Felids are the only definitive host for the parasite, while rodents, birds and humans serve as an intermediate host (McConkey et al. 2013). However, this single-celled parasite preferred to reside in the brain of its intermediate host (Berenreiterová et al. 2011). Interestingly, it has been reported that prolong *T. gondii* infection results into variety of behavioural deficits in rodents including altered locomotor activity, learning and memory impairment (Nolan 2019). Such an interplay between rodents infected with *T. gondii* parasite facilitates its transmission back to its felids definitive host, suggesting the ability of the parasite to impair the host behaviour (Abdulai-Saiku and Vyas 2017; Berdoy, Webster, and Macdonald 1995; McConkey et al. 2013; Webster 2001). Nevertheless, previous research in humans have documented a strong association between *T. gondii* infection and mental disorder such as alzheimer, suicidal attack, bipolar disorder and schizophrenia (Elsheikha, Büsselberg, and Zhu 2016; Esshili et al. 2016; Sutterland et al. 2015).

Schizophrenia is a multifaceted mental disorder with unknown aetiology (Esshili et al. 2016). This mental disorder is one of the major causes of a psychiatric condition that has ravaged human population since time immemorial with an estimated 1% of people infected with toxoplasmosis (Flegr, 2015; Yolken et al., 2009). Rodent models of *T. gondii* infection serve as a tool to investigate behavioural changes induced by the parasite. Several studies have demonstrated a wide range of subtle behavioural changes in laboratory rat and mice infected with *T. gondii* (Afonso et al., 2017; Berdoy, Webster, & Macdonald, 1995; McConkey et al., 2013; Skallová et al., 2006; Webster et al., 2013). Factors such as impaired aversion of rodent to cat odour (Berdoy, Webster, & Macdonald, 2000; Flegr et al., 2011; Kaushik, Knowles, & Webster, 2014; Webster, 2001), fear and anxiety (Evans et al., 2014; Gonzalez et al., 2007; Ihara et al., 2016; Mahmoudvand et al., 2015), deficits in learning and memory retention (Anisman & McIntyre, 2002; Bezerra et al., 2019; Daniels et al., 2015; de Bruin et al., 2001; Hodkova, Kodym, & Flegr, 2007), object recognition test (Lamberton, Donnelly, and Webster 2008), and higher activity in infected rat (Gonzalez et al., 2007; Kaushik et al., 2014; Webster, 1994). In spite of huge *T. gondii* research evaluating behaviour deficits in rodents worldwide, the effects of Malaysian strains of *T. gondii* have not been tested on rat’s behaviour (Wana et al. 2020a).

To date, research have documented views on the mechanism(s) supporting behavioural changes after *T. gondii* infection and the subsequent establishment of tissue cysts in the brain of rodents. The mechanism (s) underlying the direct effect related to the distribution, localization or tropism of *T. gondii* tissue cysts and behavioural changes has been reported (Berenreiterová et al., 2011; Daniels et al., 2015). In infected host, *T. gondii* tissue cysts were found distributed throughout the brain and the strain causing the infection may likely represent subtle behavioural deficits observed in rodents (Bezerra et al., 2019). Studies in rats have also attributed lack of specific tissue cysts tropism in the anatomical brain regions from the prefrontal cortex to occipital (Gonzalez et al. 2007) may likely explain the effect of different *T. gondii* strain.

Nevertheless, only a few studies in rodents have examined *T. gondii* tissue cysts presence and location linked to the differences in *T. gondii* strains (Bezerra et al., 2019; Kannan et al., 2010). This is the first report that tested the effects of Malaysian strains of *T. gondii* infected rat with respect to possible cognitive impairments. The study also evaluated the distribution *T. gondii* tissue cysts in the rat’s brain regions relevant to learning and memory processing.

## 2. Material and Methods

### Genetic characterization of *T. gondii* oocysts

In a previous study, 17 cat faecal samples were confirmed positive from the pet and free roaming cats in Klang Valley, Malaysia (Wana et al. 2020b). The strains typing of *T. gondii* isolates was conducted with SAG3 genetic marker which was designed previously (Grigg and Boothroyd 2001; Su, Zhang, and Dubey 2006). The genetic marker is capable to distinguished *T. gondii* genotype strains into type I, II, and III strain. The reaction mixture contains 5 μl of the DNA template, 25 μl of the Superhot Master Mix (BIORON GmbH, Germany), 0.1 μg/μl concentration of BSA, 0.2 μM (final concentration) of forward and reverse external primers (F: CAACTCTCACCATTCCACCC, R:GCGCGTTGTTAGACAAGACA) and PCR grade water. The mixture was amplified in Bio-Rad Mycylcer (Thermal Cycler PCR, USA). The initial temperature was set at 95°C for 3 min. The PCR programme was followed by 30 cycles of denaturation at 95°C for 45 s, annealing at 55°C for 1 min and extension at 72°C for 1 min. The final extension was carried out at 72°C for 5 min. The reference strain of type I, type II and type III were also prepared and run together as a positive control, while PCR mixture without DNA template was also included as negative control. The PCR product and 100 bp ladder (BIORON, GmbH, Germany) was resolved in 1.5% agarose gel stained with ethidium bromide and visualized with Bio-Rad gel doc XR (Molecular Imager, USA).

The product from the first round PCR was used as a template for the second round Nested PCR. The reaction mixture comprised of 4 μl of PCR product, 0.2 μM of forward and reverse internal primers (F: TCTTGTCGGGTGTTCACTCA, R: CACAAGGAGACCGAGAAGGA), 12.5 μl of Superhot Master Mix (BIORON GmbH, Germany) and PCR grade water. The initial denaturation was carried out at 95°C for 2 min. The subsequent steps were performed for 35 cycles with denaturation at 95°C for 45 s, annealing for all the seven internal primers at 55°C and extension at 72°C for 1.5 min. The final extension was set at 95°C for 10 min. Similarly, positive reference strains of type I, II, III and negative control from the first round were also included in the second PCR run. The PCR product and 100 bp ladder (BIORON, GmbH, Germany) were resolved in 1.5% agarose gel stained with ethidium bromide and visualized with Bio-Rad gel doc XR (Molecular Imager, USA). This was followed by restricted fragment length polymorphism (RFLP). The Nested PCR product (5 μl) was digested with NciI restriction enzyme at 37°C for 15 min (New England Biolabs, USA). The final digested DNA fragments was resolved in 2.5 % gel electrophoresis stained with ethidium bromide and visualized with Bio-Rad gel doc XR (Molecular Imager, USA) to reveal the RFLP band patterns.

### Experimental design and *T. gondii* infection

Forty-five male Wistar albino rats, four weeks old were randomly divided into five groups which contained nine rats each. The rats were assigned into five groups: two control groups (CG1 and CG2) and three experimental groups (EG1, EG2, EG3). CG1 rats received phosphate buffered saline (PBS), CG2 were injected with 0.6 mg/kg MK-801 (as a model for schizophrenia) (Wang et al. 2013), EG1, EG2, EG3 received orally 5 × 10^3^ single *T. gondii* oocysts strain of type I, type II and type III respectively. The rats were housed three rats per cage and placed under the same laboratory condition. The room was ventilated with a temperature of between ± 22°C, and 12h light-dark cycle for a period of one week to acclimatize. These animals were fed with standard commercially available rats chow with water provided *ad libitum*. All the experimental protocols were conducted according to the Animal ethics guideline and approved by the UPM Animal Care and Use Committee (UPM/IACUC/AUP-R071/2017, Appendix 1). The experimental design for the behavioural study is summarized in Figure 1. Body weights were recorded at weekly intervals during the entire period of study.

**Figure 1.**
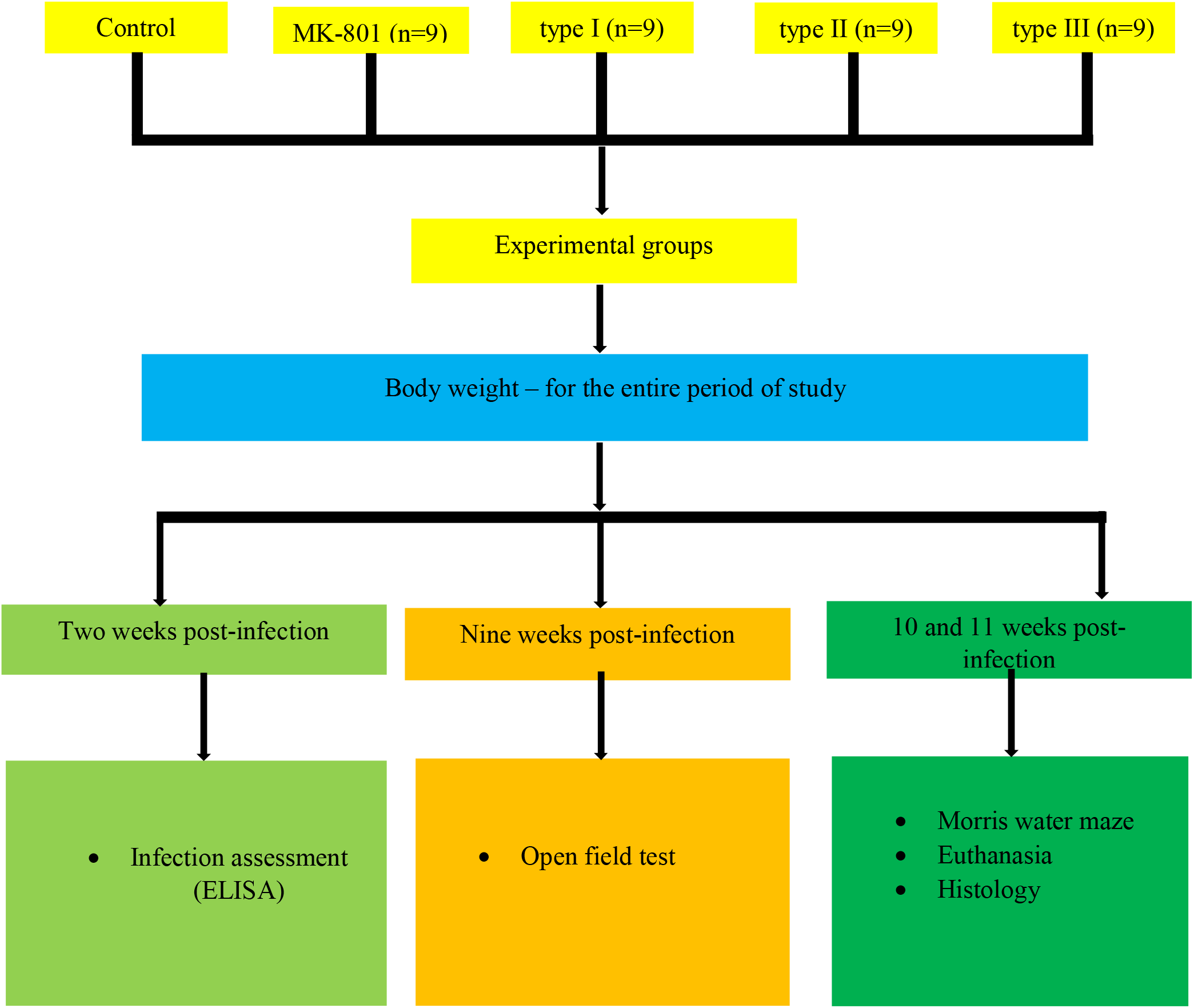
Experimental design for the behavioural study

### *T. gondii* infection determination

Blood (750 μl) samples were collected at two weeks post infection (PI) from each rat into a labelled container from the tail vein using a syringe (27G) and needle. The blood sample was immediately centrifuged at 3000 rpm for 10 mins to remove the sera and kept at −20°C until further processed. *T. gondii* antibody testing was done using the commercially available Toxoplasma IgM EIA Well (RADIM Diagnostic Pomezia, Italy) according to manufacturer’s instruction. To detect IgM of *T. gondii* antibodies, optical density (OD) calculation of the plates were performed with a microtiter plate reader (450 - 630 nm, DYNEX, MRX).

### Behavioural test at nine weeks post infection

The behavioural test of open field test (OFT) were carried out at nine weeks PI, a period which is known by *T. gondii* to established chronic infection in the rat with tissue cysts formation (Vyas et al., 2007; Vyas, Kim, & Sapolsky, 2007).

### Open field test

The OFT is a measure of the locomotor and exploratory activity of the rats when they are placed into a new environment (Chiroma et al., 2018; Bezerra et al., 2012). The OFT apparatus is a squared box made up of Plexiglas that had closed base an open top (75 cm × 75 cm base, 40 cm height). The base of the box was equally divided with parallel vertical and horizontal lines into 25 smaller square units measuring 15 cm × 15 cm. The rats were moved into the behaviour room and allowed for 10 mins to acclimatize to the room set up. Each rat was placed at the centre of the box and allowed to explore the open field arena for 5 mins (Bezerra et al., 2012). Logitech camera was fixed to the ceiling directly facing the maze which is connected to ANY-Maze video tracking software for recording. At the end of the time allotted, the rat was removed from the box and returned back to its cage. Between each rat test, 70% ethanol was used to clean the box wiped with sterile water and allowed to dry for 2 min in order to remove the odour. The number of lines crossed by the rats were recorded (Webster et al., 2013).

### Behavioural test at 10th and 11th weeks post-infection

The behavioural test of Morris water maze (reference memory acquisition and reversal learning task) were carried out at 10 and 11 weeks PI.

### Morris water maze

#### Reference memory acquisition

The Morris water maze (MWM) is a task that was widely used to evaluate spatial learning and memory paradigm in cognitive impairment of rodents (Anisman and McIntyre 2002; Daniels et al. 2015; Dass et al. 2011; Mitraa, Sapolskyb, and Vyasa 2012; Vorhees and Williams 2006; Chiroma et al., 2019). The reference memory acquisition (RMA) of MWM test consisted of a black circular tank that measured 210 cm in diameter, 80 cm height as shown in Appendix 2. The tank is filled to the half with opaque water to reduce visibility. The temperature of the water and the test room was maintained at 19-22°C. The escape platform made up of plastic (10 cm × 10 cm) was submerged in the water approximately 2 cm below the surface. Further, the tank was divided into four quadrants as N, S, E and W with each having specific distal cues. The escape platform was fixed at the N quadrant for the reference memory acquisition for all the testing days. All experiments were carried out with indirect dim light in the test room to avoid interfering with the tracking video (Vorhees and Williams 2006). The design of the camera and video tracking is similar to OFT above. Rats were evaluated on the acquisition of reference memory starting on the 10 weeks PI and were tested for 5 consecutive days. This constituted the trial phase of the task. On each testing day, each of the rats was placed in four different random positions to locate the hidden escape platform in the tank. At the end of 60 s for each trial, rats that failed to locate the hidden escape platform was guided gently to it. Each rat was allowed to rest on the escape platform for 15 s to familiarize with the environment using distal cues as a guide, after which it was removed, clean and dried, then returned to its home cage before the next rat test. On the 6 days, the hidden escape platform was removed and each rat was gently released into the water to locate the quadrant where it was positioned (Vorhees and Williams 2006). This is called the probe test and performed once 24 hrs at the end of the trial test. The test was conducted for a period of 30 s. The time spent at the target quadrant, as well as latency to reach the target quadrant were recorded as a measure of reference memory retention.

#### Reversal learning task

At the 11 weeks PI, all rats were additionally tested on MWM for the reversal learning task (RLT). The position of the platform was changed to S quadrant opposite to the initial location. The same procedure was followed for the trial and probe test as described during RMA above.

#### Haematoxylin and eosin staining of rat brain sections

Three rats per each experimental group were randomly selected from the five groups containing six rats at the completion of the behavioral test at 11 weeks post infection. The rats were humanely euthanised through decapitation (Mitra, Sapolsky, and Vyas 2013), and the whole brain was removed and placed in ice cold saline for cleaning purpose. The brain samples were removed and placed in a clean container that has 10% paraformaldehyde for preservation. The whole rat brain was fixed for three days in 10% paraformaldehyde. Thereafter, the whole rat brain was processed and subsequently fixed with paraffin wax. Using microtome machine, the block of whole rat brain tissue was cut at coronal sections from the prefrontal cortex to the occipitals. The sectioned tissues produced continues ribbons after 5 μm (Gatkowska et al., 2012; Gulinello et al., 2010) and placed on a labelled slide. Finally, slides were stained with haematoxylin and eosin and mounted with dibutyl phthalate xylene (DPX), then covered with a coverslip. The presence of cysts, distribution and scoring were recorded with a guide of histologist under 100x and 400x magnification with a microscope (Olympus BX51TRF-CCD).

#### Scoring method of *T. gondii* tissue cysts

Scoring of *T. gondii* tissue cysts in the brain of rats were performed through the guide of histologist which followed the methods described by Berenreiterova et al., (2011). The average number of tissue cysts were recorded based on anatomical positions of three rats per each experimental group (n=3). The size of the tissue cysts range between 10 and 100 μm and were found solitary or in a group of two to six. Majority of the tissue cysts (70%) were found solitary, while few (30%) in groups.

### Statistical analysis

GraphPad Prism (version 7.0) software (GraphPad Software, San Diego, CA, USA) was used to analysed body weight, locomotor activity, probe test of Morris water maze, tissue cyst distribution and brain sections through one way analysis of variance (ANOVA) followed by Tukey’s post hoc test. Data generated from trial phase of Morris water maze were analyzed through two way ANOVA followed by Tukey’s post hoc test. A comparison was made between the experimental groups with a value of P < 0.05.

## 3. Results

### *T. gondii* strains in Malaysia from cat faeces

The 17 *T. gondii*-positive DNA samples directly from cat faeces, all were successfully genotyped at SAG3 locus. Overall, results revealed Malaysian strains of T. gondii are clonal with predominant type I. Further, the results clearly shows 10 faecal samples belong to type I, 4 type II and only 1 type III. The RFLP band pattern is illustrated in Figure 2.

**Figure 2.**
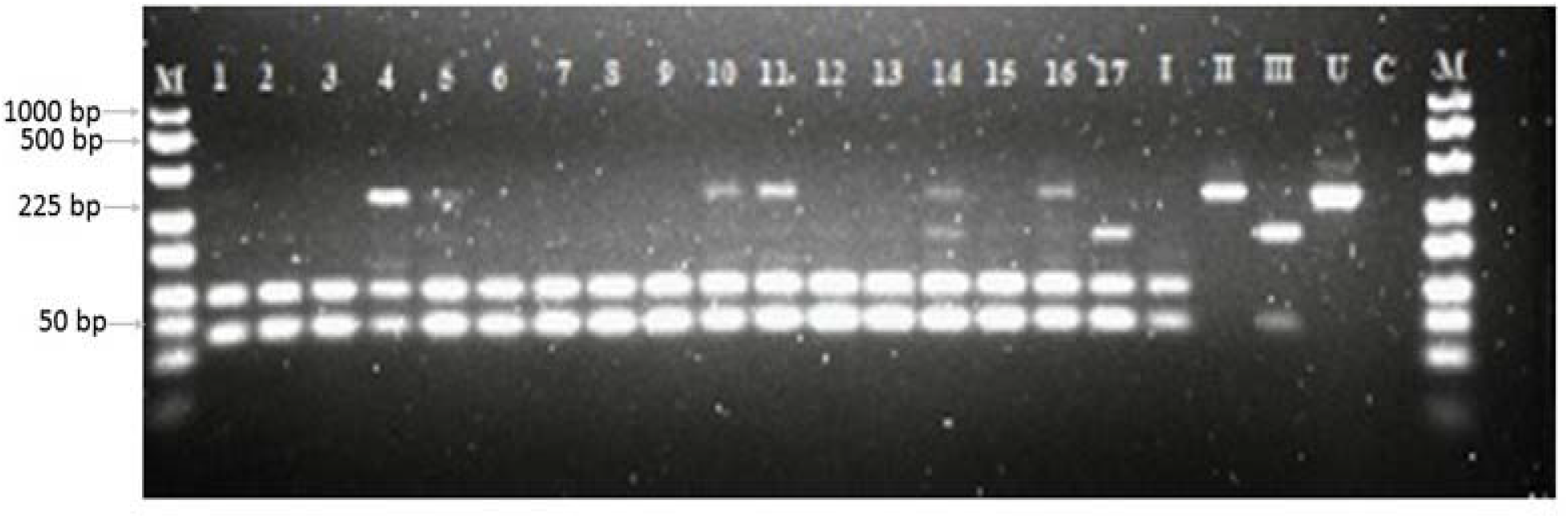
Genotypic characterization of *T. gondii* strain using genetic marker, SAG3. The RFLP product and undigested product was resolved in 2.5% agarose stained with Redsafe. Lane M: 100 bp DNA molecular ladder (GeneRuler, Thermo Scientific); Lanes 1-17: *T. gondii* isolates from Malaysia; Lane I, II and III reference strains of type I (RH), type II (PRU) and type III (VEG) respectively. Lane U; undigested product, lane NEG; negative control (PCR reaction mixture without DNA).

### Serological analysis

The serological test of IgM found at least 6, 7 and 6 rats in EG1, EG2 and EG3 respectively *T. gondii* infected groups were positive, while control groups of rats CG1 and CG2 were negative. Further, the experimental groups were normalized to have six rats per group. Only rats that were positive with *T. gondii* antibodies in EG1, EG2 and EG3 proceeded to the behavioural tests and histological analysis.

### Body weight

The body-weight of the MK-801 induced model group of schizophrenia, *T. gondii* infected groups and control group of the rat were comparable at the end of 8 weeks post infection before the commencement of behavioural test (Figure 3).

**Figure 3.**
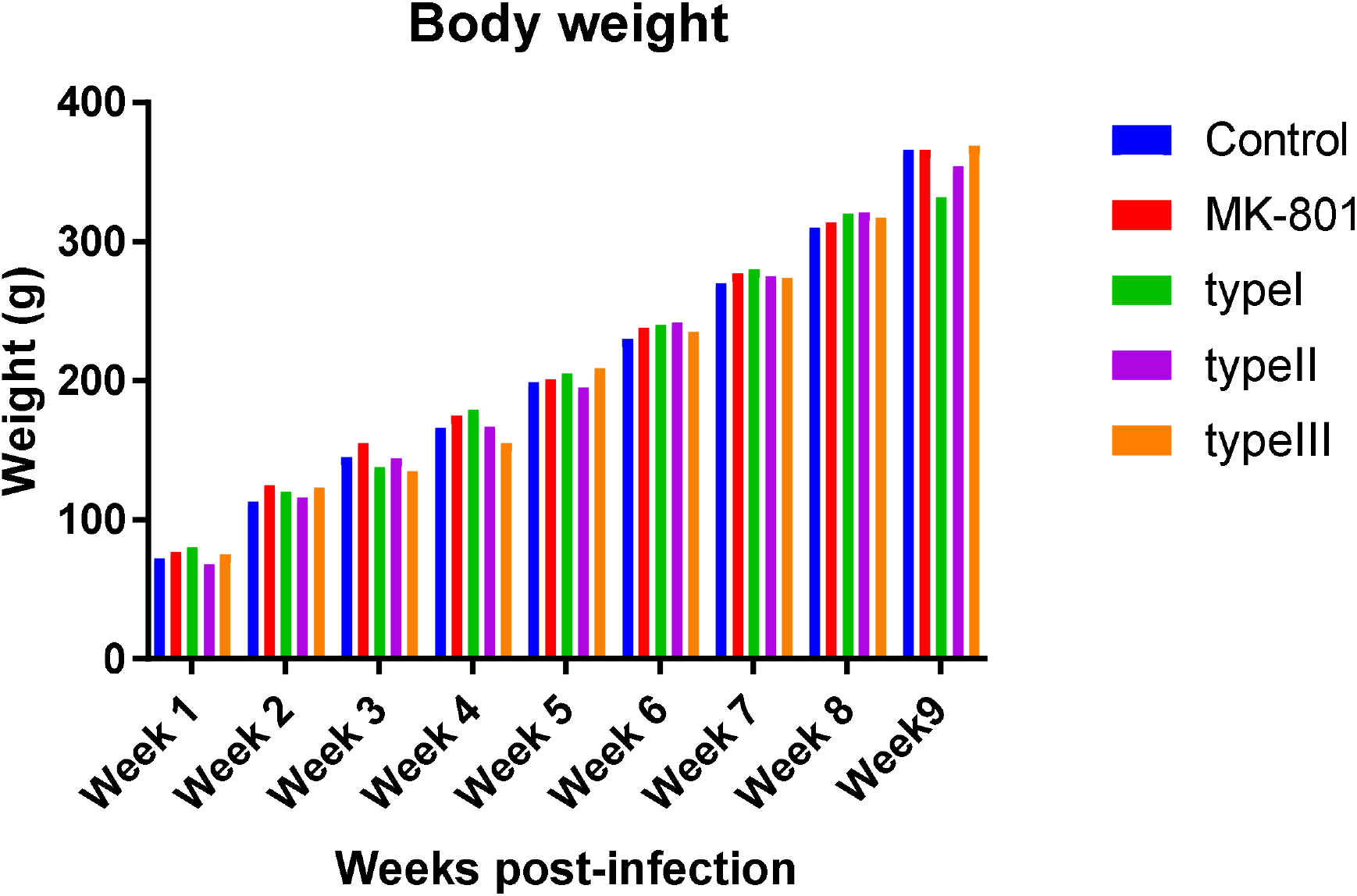
Body weight of Wistar rats administered with MK-801 group, infected with various genotype strain of *T. gondii* groups compared with the control group of rat post-infection. Significant differences among body weight were analysed through one-way ANOVA followed by Tukey’s post hoc test, but there was comparable body weight between the groups of rats throughout the entire period of the study. Data were expressed as average (n=6) body weight for each group.

### Locomotor activity

There is increased in locomotor activity in MK-801 and *T. gondii* infected experimental groups of rats compared with the control group of rats. Further, the OFT did reveal a statistical significance in locomotor activity between MK-801 and *T. gondii* infected group compared to the control group of rats (Figure 4) and (Figure 5). Several *T. gondii* infected rats and MK-801 induced rats exhibited circular activity along the edge of the box arena which effectively increases the mean number of line crosses, whereas the control rats showed less mean number of line crosses. However, data generated did not show defects in locomotor activity in all experimental groups.

**Figure 4.**
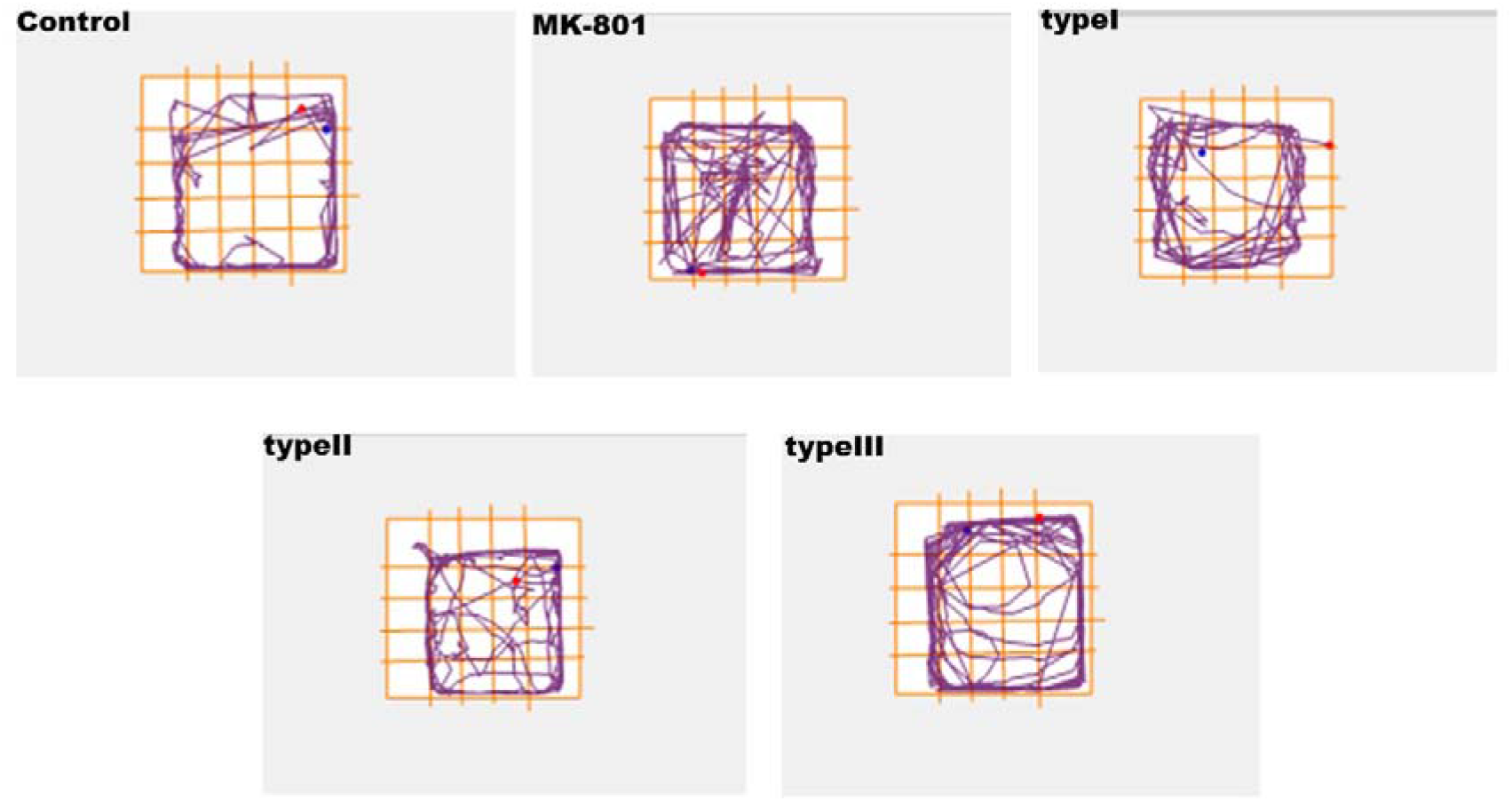
Representatives of track plots of locomotor activity of control, MK-801 and *T. gondii* infected experimental groups of rats in OFT. A represents the control rat group, B represents the MK-801 model group of rat, while C, D and F represent the *T. gondii* infected groups of rats from type I, II and III respectively (n=6).

**Figure 5.**
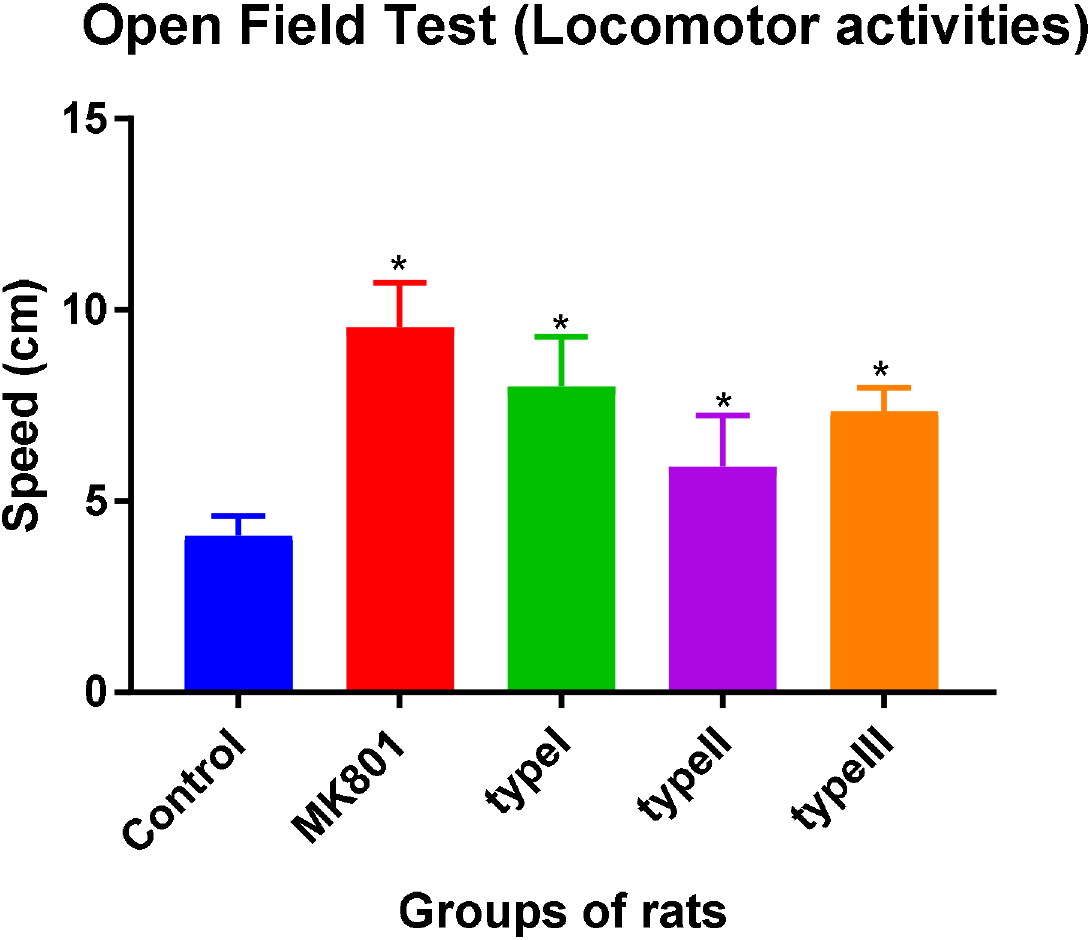
Open field activity of *T. gondii* infected and MK-801 groups of rats on locomotor activity. There is increased in locomotor activity in both *T. gondii* infected and MK-801 groups compared with the control group using one way ANOVA. Further, Turkey’s post hoc test show a statistically significant difference between MK-801 and *T. gondii* infected experimental group

### Spatial learning and memory retention

The result revealed a reference to spatial learning and memory retention in *T. gondii* infected rats using MWM. The movement pattern during the spatial learning phase is depicted in Figure 6. The escape latency to reach the hidden platform was consecutively reduced in a 5 days trial for all the experimental groups (Figure 7A). The data analysis using 2-way repeated measure of ANOVA indicated that there is no difference in the time taken by the rats to locate the hidden platform (P < 0.001), as all groups of rat, perform equally. Further spatial memory retention, 24 hrs after the last spatial learning was carried out, a probe test (Figure 7B). The parameter measured was the time spent in the target quadrant when the hidden platform was removed as an indicator of memory retention. However, one way ANOVA revealed a statistical significance difference [F_(4, 19)_ = 11.55, P < 0.001] between the various experimental groups of rats. This was followed by Tukey’s post hoc test which revealed a statistical significance difference between MK-801 induced (3.3±2.3) and *T. gondii* infected of type I (6.1±2.9), type II (7.3±0.7) and type III (5.1±2.0) groups of rats when compared to the control group of rats (13.63±3.5). This finding indicated a memory deficit and poor search strategy in MK-801 induced and *T. gondii* infected rat groups to locate the initial location of the hidden platform as a target quadrant. To further test for spatial learning and memory retention, a reversal learning process was conducted. This is similar to the reference test performed earlier, except that the hidden platform was placed in a new quadrant directly opposite the first location. Results obtained when rats were forced to learn and retain the memory of new location followed the same pattern of reference spatial learning and memory retention test. This indicated a substantial learning and memory impairment in MK-801 and *T. gondii* infected groups of rat.

**Figure 6.**
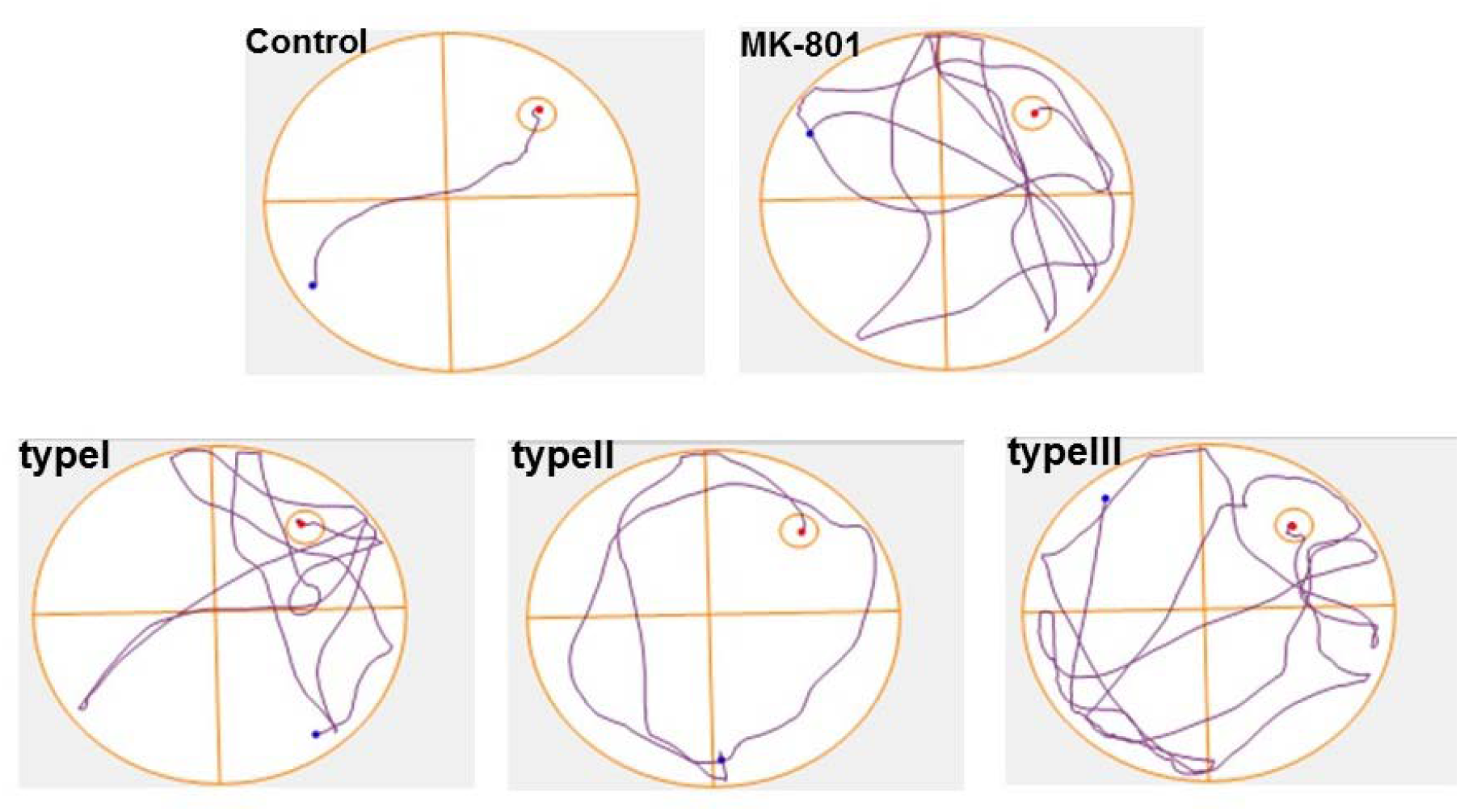
Representative images of track plots of spatial learning and memory retention of control, MK-801 and *T. gondii* infected experimental groups of rats in MWM. The control groups comprised; CG1, control group, inoculated with PBS. The CG2, administered with (+) MK-801, a rat model of schizophrenia. The experimental groups, EG1 infected with type I strain; EG2 infected with type II strain; EG3, infected with type III strain.

**Figure 7.**
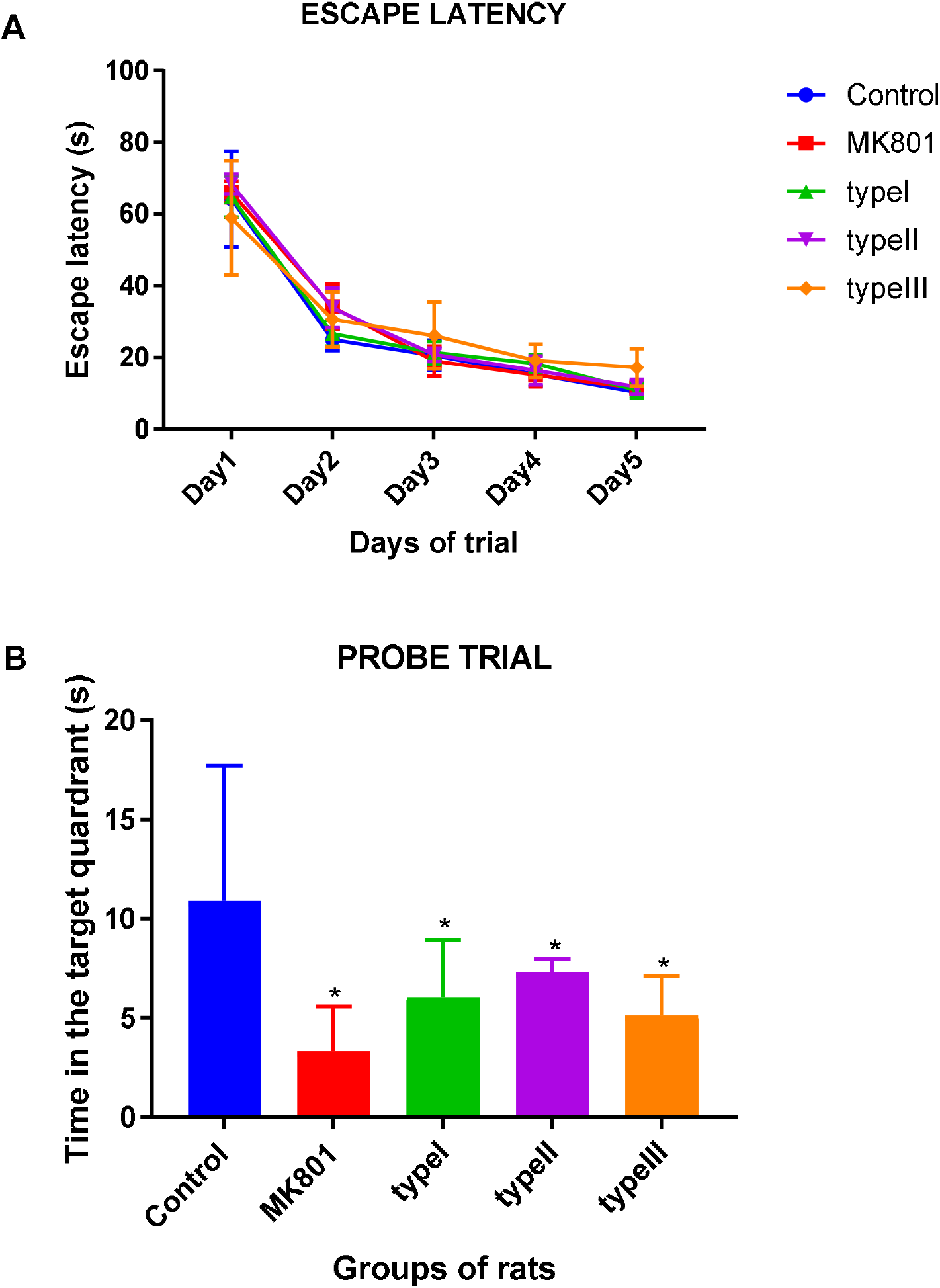
Evaluation of spatial learning and memory of rats in MWM. A) Escape latency (s) during 5 days trials to locate the hidden platform as part of the learning phase in the Morris water maze. B) The probe trial conducted after the trial phase when the hidden platform is removed. Rats were allowed to search for the platform in the target quadrant. The control groups comprised; CG1, control group inoculated with PBS. The CG2, administered with (+) MK-801, a rat model of schizophrenia. The experimental group, EG1 infected with type I strain; EG2, infected with type II strain; EG3, infected with type III strain. Data were presented as ± SEM for (n=6) rats per group. *P < 0.05 vs control group. Error bars indicate SEM (n=6).

### The distribution of the *T. gondii* tissue cysts in the brain of infected rats

Figure 8 shows a photomicrograph of rat brain section of the control groups induced with PBS, MK-801 administered group and *T. gondii* infected groups of rats. The control group is healthy rats that have a normal brain section. Severe lesions were seen all over the brain of rats administered with MK-801 0.6 mg/kg to mimic schizophrenia as a model group. Alternatively, a group of rat infected with *T. gondii* genotype strain of type I, type II and type III have tissue cysts present all over the brain without tropism.

**Figure 8.**
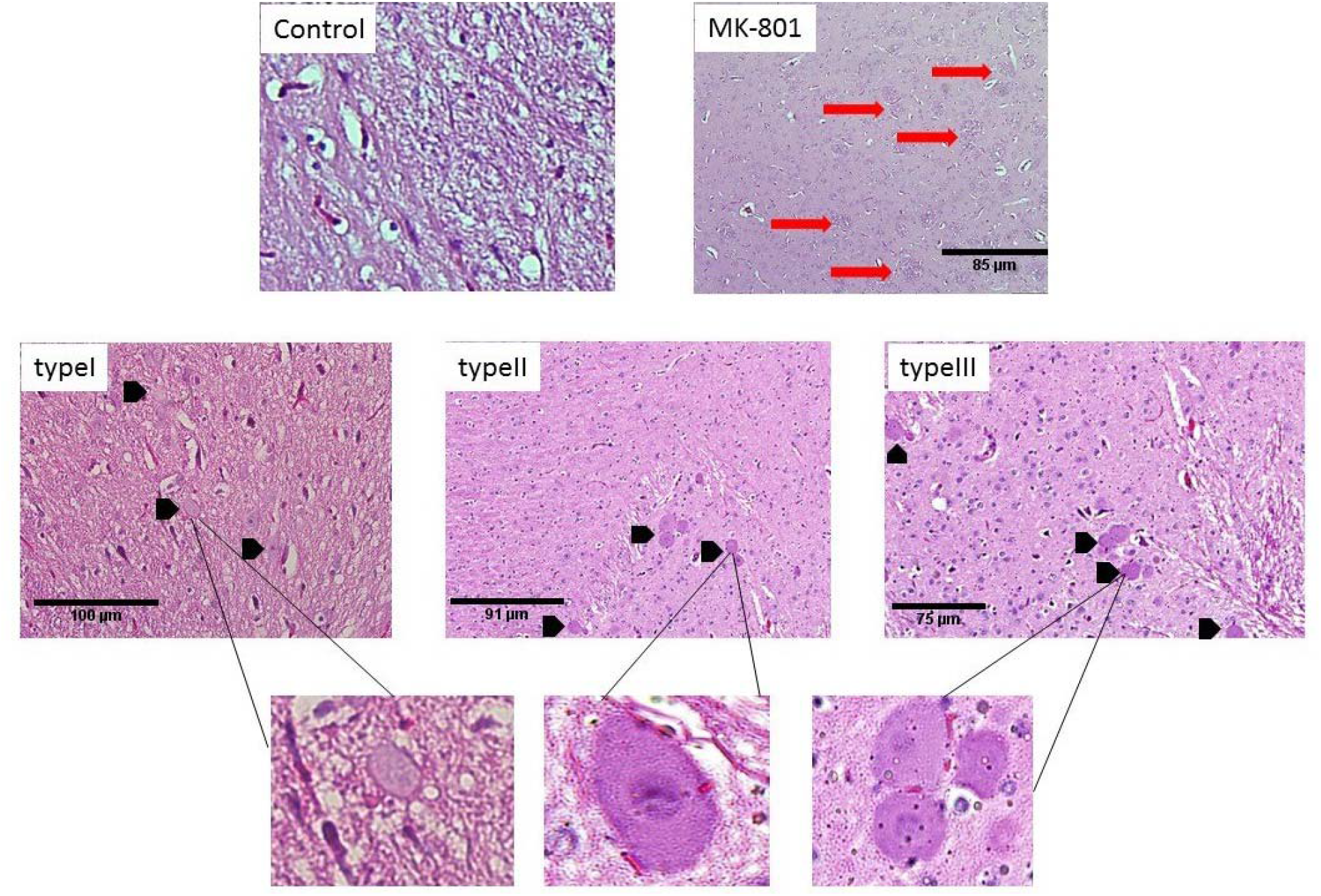
Micrographs of Haematoxylin & Eosin stained rat brain sections from five experimental groups. The control experimental group of rat shows a normal brain section (Magnification of 100x). The schizophrenia model experimental group of rat injected with MK-801 shows lesions pointed with red arrows all over the brain section (Magnification of 100x). The type I, type II and type III are the three experimental groups of rats infected with type I, II and III *T. gondii* genotype strains showing tissue cysts indicated with arrow-head (Magnification of 100x). At the bottom are enlarged *T. gondii* tissue cysts from type I, II & III present in infected rat brain sections indicated with black arrows. Scale bar were analyzed through ImageJ (version FIJI 1.46r, National Institute of Health) software.

### *T. gondii* tissue cysts distribution in the brain sections of rats based on anatomical position

Figure 9 shows the distribution and number of *T. gondii* tissue cyst were found in seven distinct brain section which include cerebral cortex, corpus callosum, nucleus accumbens, hypothalamus, thalamus, amygdala and caudate putamen. The three *T. gondii-infected* experimental groups of typeI, type II and type III contain the tissue cysts. One way ANOVA showed a statistical significant difference in the number of tissue cysts among the experimental groups of rats infected with *T. gondii*. Further, Tukey’s *post hoc* test revealed no statistical difference between the number of tissue recorded between the brain sections and the three *T. gondii* (type I, II and III) experimental groups. In addition, tissue cysts were not seen in the prefrontal cortex, hippocampus and cerebellum in all the five experimental groups of rats. Similarly, the other two experimental groups of control and MK-801 were devoid of *T. gondii* tissue cyst.

**Figure 9.**
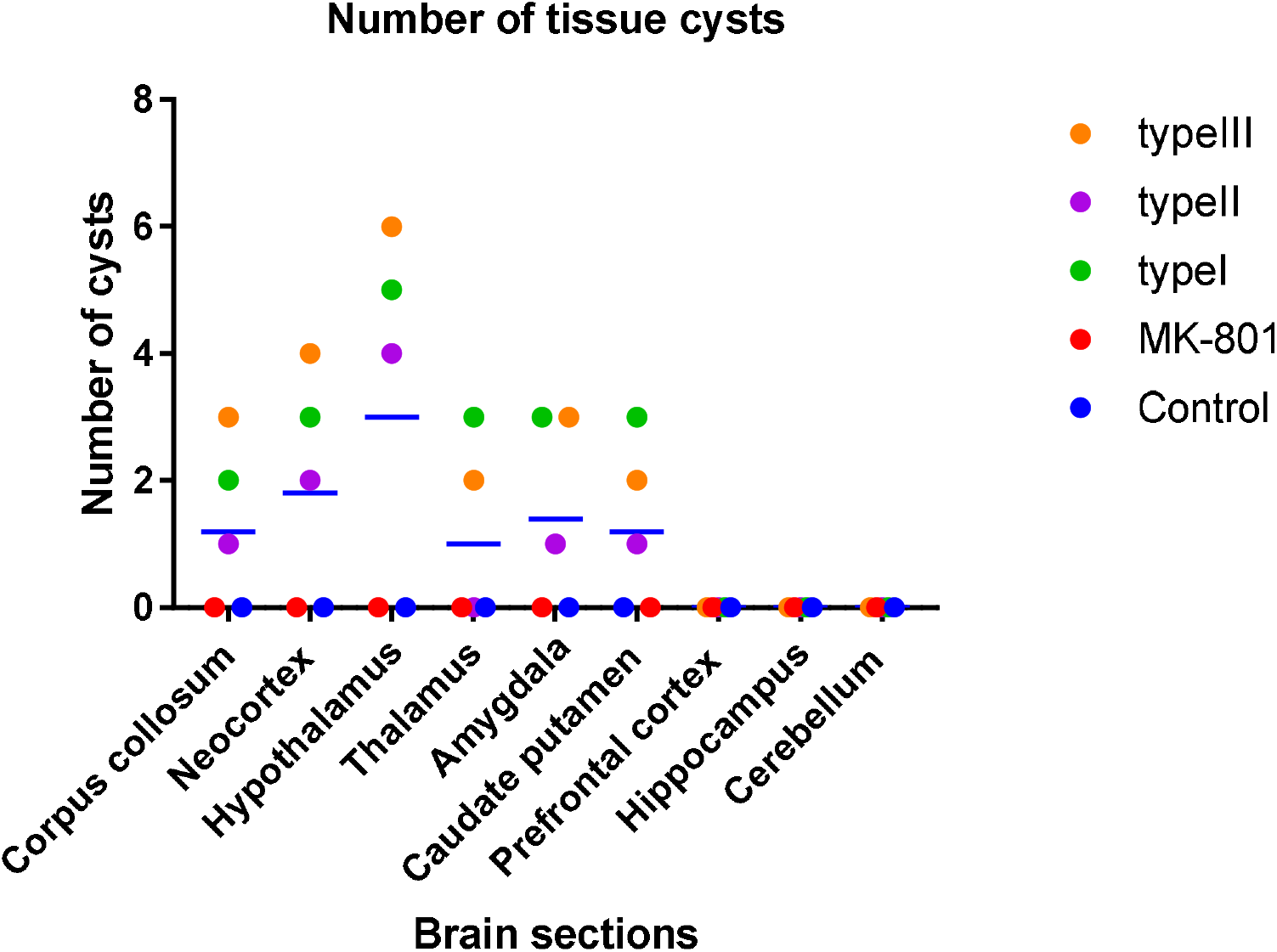
Analysis of *T. gondii* tissue cyst in the brains of infected rats. The ordinate indicates the number of tissue cyst in each brain sections. Rat whole brain was collected at 12 weeks postinfection. The circle data points represent data for one rat, and bars indicate the average value of three infected rats and their particular brain section (*T. gondii*-infected rats, n=3). Statistically, no significant differences were found between the numbers of *T. gondii* tissue cyst among the infected groups.

**Figure 10.**
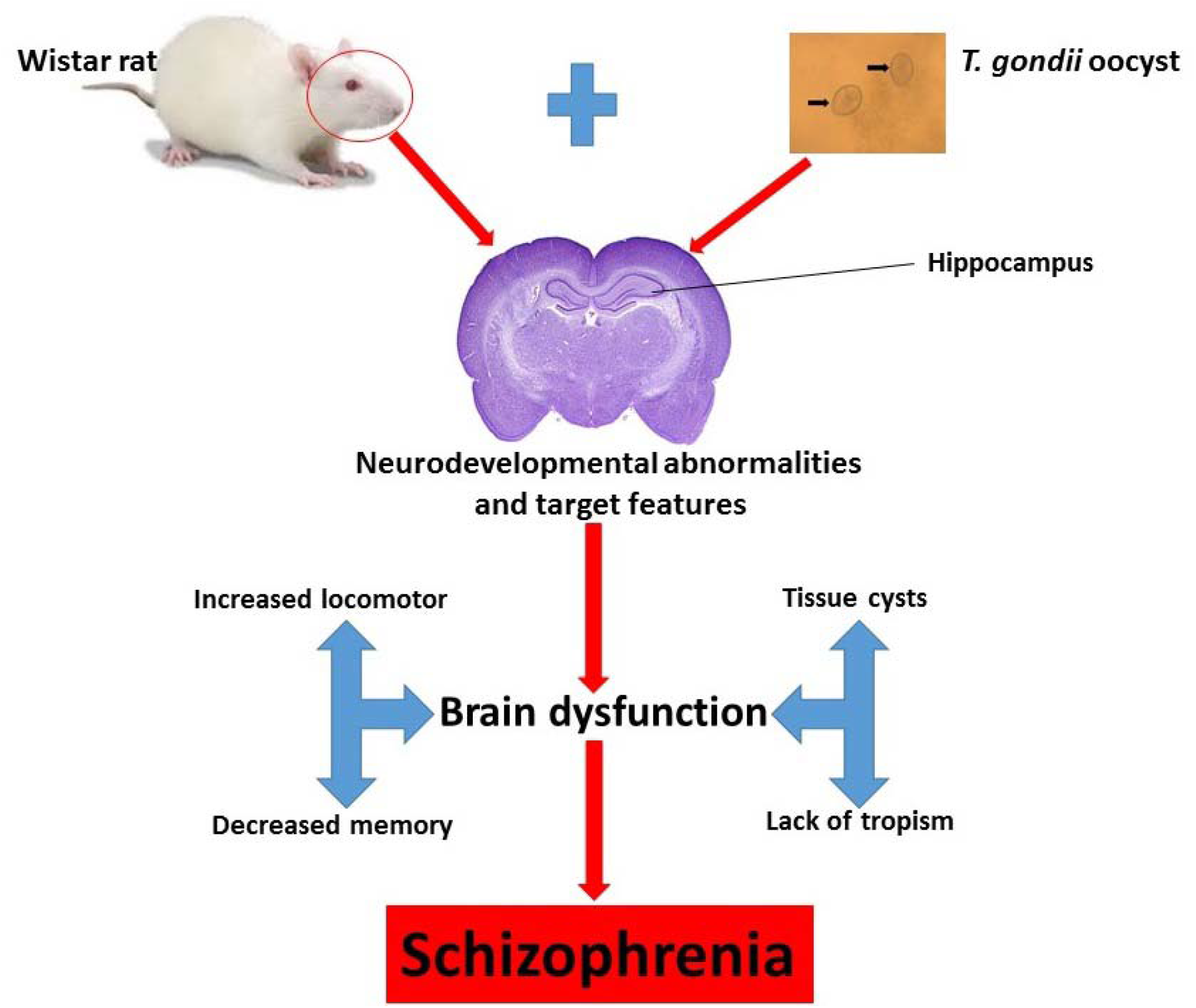
Proposed mechanism of *T. gondii* induced neurodevelopmental abnormalities and cognitive deficits in rats. Inoculation of *T. gondii* oocysts to Wistar rats resulted to abnormal function of principal brain regions such as hippocampus, amygdala and hypothalamus that lead to increase in locomotor activity, loss of memory retention and presence of tissue cysts without tropism to a particular region in the brain of rats. These behavioural changes mimic the cognitive deficits observed in schizophrenia.

## 4. Discussion

*T. gondii* is a unicellular obligate protozoan parasite that can localize mainly in the brain of its intermediate host which remain as a chronic/latent infection throughout the entire life of the host causing behaviour deficits (Tyebji et al., 2019).

The increased in locomotor activity in both *T. gondii* infected and MK-801 experimental groups of rats found in the present study (Figure 5 and Figure 6) which is consistent with previous findings in mice infected with different strains of *T. gondii* (Afonso et al., 2012; Hodkova et al., 2007; Kannan et al., 2010; Xiao et al., 2012). In contrast, other studies in mice infected with *T. gondii* have reported reduced locomotor activity in an open field test (Gulinello et al., 2010; Xiao et al., 2016). However, lack of changes in locomotor activity was also found in mice (Kannan et al., 2010; Xiao et al., 2012) and rats (Gonzalez et al. 2007; Vyas, Kim, Giacomini, et al. 2007) infected with *T. gondii*. These mixed findings were typically associated with differences in the choice of rodent animal species and their sex, *T. gondii* strain and behavioural apparatus (Tyebji, et al., 2019), suggesting that *T. gondii* effect on neuronal cells and test for locomotor activity in an open field may produce variable outcomes.

In the present study, MK-801 induced and *T. gondii* infected groups of the rat were found to have intact spatial learning potentials in a MWM task, but memory retention was impaired. While MK-801 and *T. gondii* infected group learned the location of the hidden platform compared to the control group, they suffered memory loss during probe trials (Figure 7B). This is line with previous studies that reported a lack of deficits in learning task of the MWM (Vyas et al., 2007). However, Daniels et al. (2015) further demonstrated memory deficits in rats during probe trials of MWM which is in agreement with the findings of the present study. Conversely, the present study has found no obvious variation between the behavioural effects of type I, type II and type III *T. gondii* strain. In contrast, spatial memory impairment in mice was associated with ME49 rather than PRU strain (Kannan et al., 2010). In the other study in mice (Jung et al. 2012) they have found no obvious learning and memory deficit. Intriguingly, only a few studies have documented learning and memory alteration in laboratory rodents infected with *T. gondii* using MWM, which largely utilized the Y-maze (Kannan et al., 2010). In addition, behavioural studies in *T. gondii* infected animals have focused more in mice, which are highly affected with acute infection and less likely to develop chronic infection (Kannan et al., 2010; Worth, Lymbery, & Thompson, 2013). These discrepancies in the previous findings on the impact of *T. gondii* on behaviour changes could likely result from the choice of rodent species, apparatus used for the test and time PI (Tyebji et al., 2019; Worth et al., 2013). The changes observed mostly in *T. gondii* infection that has to do with behaviour changes is related to chronic infection (Tyebji et al., 2019) rather than acute infection. Whereas, acute infection involves a short duration of time, possibly within one month after initial infection (Vyas et al., 2007), chronic infection ensured long term establishment of the parasite in the brain after acute infection has resolved to cause behaviour alteration (Kannan & Pletnikov, 2012; Vyas et al., 2007). Since chronic infection is connected with deficits in learning and memory consequences, the present study examined *T. gondii* infected rats at 10 and 11 weeks PI. In this study, the duration of 10 to 11 weeks PI were found to impair memory retention of rats infected with *T. gondii* which could provide a subtle link to schizophrenia. Similarly, Kannan et al. (2012) reviewed literature found 10 weeks and above post infection when most behaviour deficits were recorded in *T. gondii* infected rodents. This issue of time factor after exposure to *T. gondii* infection could be directly responsible for brain damage and thus explain behavioural deficits.

The distribution of *T. gondii* tissue cysts in the brain of rodents have varied among studies, this could be possible because of different experimental designs, animal species, *T. gondii* strain and dose of inoculation which may have affected the outcomes (Bezerra et al., 2019; Kannan et al., 2010; Vyas, Kim, Giacomini, et al., 2007). Further, the present result (Figure 8) showed that no specific location of tissue cysts was observed, indicating a lack of cyst tropism. This was supported by previous findings (Afonso et al., 2012; Daniels et al., 2015; Gatkowska, Wieczorek, Dziadek, Dzitko, & Dlugonska, 2012b; Ihara et al., 2016) suggesting that location of tissue cyst in brain section is an opportunity rather than preference infection. Here also in this study, we reported strain-dependent distribution of *T. gondii* in the brain of infected rats. This is consistent with the previous studies that reported the distribution of *T. gondii* tissue cysts in the brain using different strains (Bezerra et al., 2019; Kannan et al., 2010; Vyas et al., 2007). However, in a recent study Bezzera et al. (2019) have found a significant increase in parasite load of VEG strain compared with ME-49. In contrast, this study did not found a significant difference in the number of tissue cysts in rats infected with either type I, type II or type III *T. gondii* strain in the brain sections. Several *T. gondii* strain have been shown to exhibit different virulent factors with type I more virulent in mice (Saraf et al., 2017), preference of intermediate host with type I commonly isolated from HIV patients (Khan et al., 2005) which is the cause of seroconversion involved in brain toxoplasma encephalitis (Djurković-Djaković et al. 2006). Therefore, this study suggests that *T. gondii* infection by different strain in healthy rat did not differ in tissue cysts distribution.

The present study, have found the presence of *T. gondii* tissue cysts in the brain sections of infected rats (Figure 9). Several reports have also documented the presence of tissue cysts in various anatomical brain regions of either mice (Ihara et al. 2016; Mahmoudvand et al. 2015; Parlog et al. 2014; Suzuki et al. 2010) or rats (Daniels et al., 2015; Vyas et al., 2007) infected with *T. gondii* cysts. However, this is the first report that showed the presence of *T. gondii* tissue cysts in the brain of rats infected with Malaysian strain of *T. gondii* oocysts. Consistent with previous findings in rats (Berenreiterová et al., 2011; Gonzalez et al., 2007; Vyas et al., 2007), the current study found tissue cysts distributed in some distinct rat’s brain region such as cerebral cortex, corpus callosum, nucleus accumbens, hypothalamus, thalamus, amygdala and caudate putamen, except that no tissue cysts were found in the prefrontal cortex, hippocampus and cerebellum (Figure 9). Also as noted by Daniels et al. (2015), no tissue cysts were detected in the hippocampus of infected rats. Meanwhile, the findings of tissue cysts in some brain sections may likely affect other parts as demonstrated by nucleus accumbens which connect amygdala and hippocampus (Daniels et al. 2015; Tan et al. 2015).

Interestingly, previous studies have reported presence of *T. gondii* tissue cysts in the hippocampus (Afonso et al., 2012; Ihara et al., 2016; Vyas et al., 2007), prefrontal cortex (Vyas, Kim, Giacomini, Boothroyd, & Sapolsky, 2007) and cerebellum (Ihara et al. 2016) of infected rodents.

Since there is lack of define location of *T. gondii* tissue cysts in the brain region that describe basic characteristics of behavioural change, specific types and magnitude of the behaviour change may vary due to area where the tissue cysts develop in the infected brain (Berenreiterová et al., 2011; Daniels et al., 2015). The hypothalamus, caudate putamen and amygdala habour *T. gondii* tissue cysts and this may likely be the reason why subtle behavioural changes were recorded in the locomotor activity, fatal feline attraction and anxiety-like behaviour. In line with the absence of *T. gondii* tissue cysts in the hippocampus, the present study observed behavioural deficits to rats learning and memory retention which may partly explain the role of the hippocampus, but may also likely results from broad spectrum alterations of information that connect functional areas of the brain. In addition, tissue cysts were found in the cerebellum and amygdala that connect with the hippocampus and are associated with learning and memory processing. Likewise, the finding of tissue cysts in the amygdala is consistent with the previous report (Berenreiterová et al., 2011; Gatkowska et al., 2012; Vyas, et al., 2007). The amygdala plays an important role in regulating cognitive and behavioural changes in memory retention, including learning processing (Maren 2001). Additionally, navigational patterned (Redish and Touretzky 1998), and changes in the amygdala was observed to cause behavior deficits in Morris water maze (Anisman and McIntyre 2002; Naghdi, Oryan, and Etemadi 2003). Other brain sections involved in memory retention, movement of the limbs, cognitive performance were found with *T. gondii* tissue cysts, including the corpus callosum which connects the two hemispheres of the brain and have direct contact with the hippocampus.

The presence of tissue cysts and its direct impact on some brain section is not the only way in which *T. gondii* can alter brain function and facilitate transmission to the cat. In addition, tissue cyst densities among the brain regions did not differ statistically where there is no significant difference between the numbers of tissue cyst recorded in each brain section. Berenreiterova et al. (2011) determined that distribution of *T. gondii* tissue cysts in the brain could be as a result of changes associated with crosses along the blood-brain-barrier (Lambert et al., 2011; Unno et al., 2008). In a previous study by Evans et al. (2014), they reported that the different behaviour phenotype observed in the infected rat is the result of influence associated with tissue cysts distribution. Although we were limited with a small number (n=3) of rats to strengthen the present findings, the histological data seem closely consistent with the behavioural changes in rats that may likely results into cognitive deficits if human were infected.

## 5.0 Conclusion

The results in this study demonstrated a possible risk to develop schizophrenia after infection with the Malaysian strain of *T. gondii*. The findings may likely provide valuable information to curb the spread of mental illness in Malaysia through prevention and control of *T. gondii* infection.

Further work is still required to check for more behavioural paradigm possibly affected and the mechanism of action of this parasite. In conclusion, rats infected with Malaysian strains of *T. gondii* showed increase in locomotor activity, poor memory retention in CG2, EG1, EG2 and EG3 compared to CG1. In addition, infection with Malaysian strain of *T. gondii* caused tissue cysts in some brain regions without distinct tropism towards a particular region. These pattern of unequal distribution of tissue cysts or absence in some brain regions may likely be due to movement along the blood-brain-barrier or blood flow. Thus, the presence of tissue cysts in corpus callosum, neocortex, hypothalamus, thalamus, caudate putamen, cerebellum and chiefly amygdala have supported the findings of behaviour deficits observed in rats and can likely occur in human if infected with *T. gondii*. These alteration in behaviour deficits observed were not strained dependent and subtle changes produced by *T. gondii* infected rats may mimic those in MK-801 model rat group suggesting schizophrenia manner. Therefore, more focus on environmental factors (*T. gondii*) is needed by policy and decision makers to curb the damaging effect of mental ill health and brain dysfunction. Thus, molecular studies are required to elucidate the probable pathways of indirect effect of *T. gondii* in rat infection.

## Author Contributions

Conceptualization of ideas, Mohamad A. M. M, Malaika W. and Roslaini AM.; Article preparation, Wana M. N, Sharif A. A, Ashraf A. Alapid, and Tijjani M.; Reviewing the initial draft and editing, Samaila Musa Chiroma, Ngah Z. U, Shariza N and Rusliza B, Final approval, all authors.

## Acknowledgment

The authors wish to extend our appreciation to Prof. Johnson Stanlas of the Department of Pharmacology, Faculty of Medicine and Health Sciences, Universiti Putra Malaysia for allowing us to use his tank for behavioural studies

---

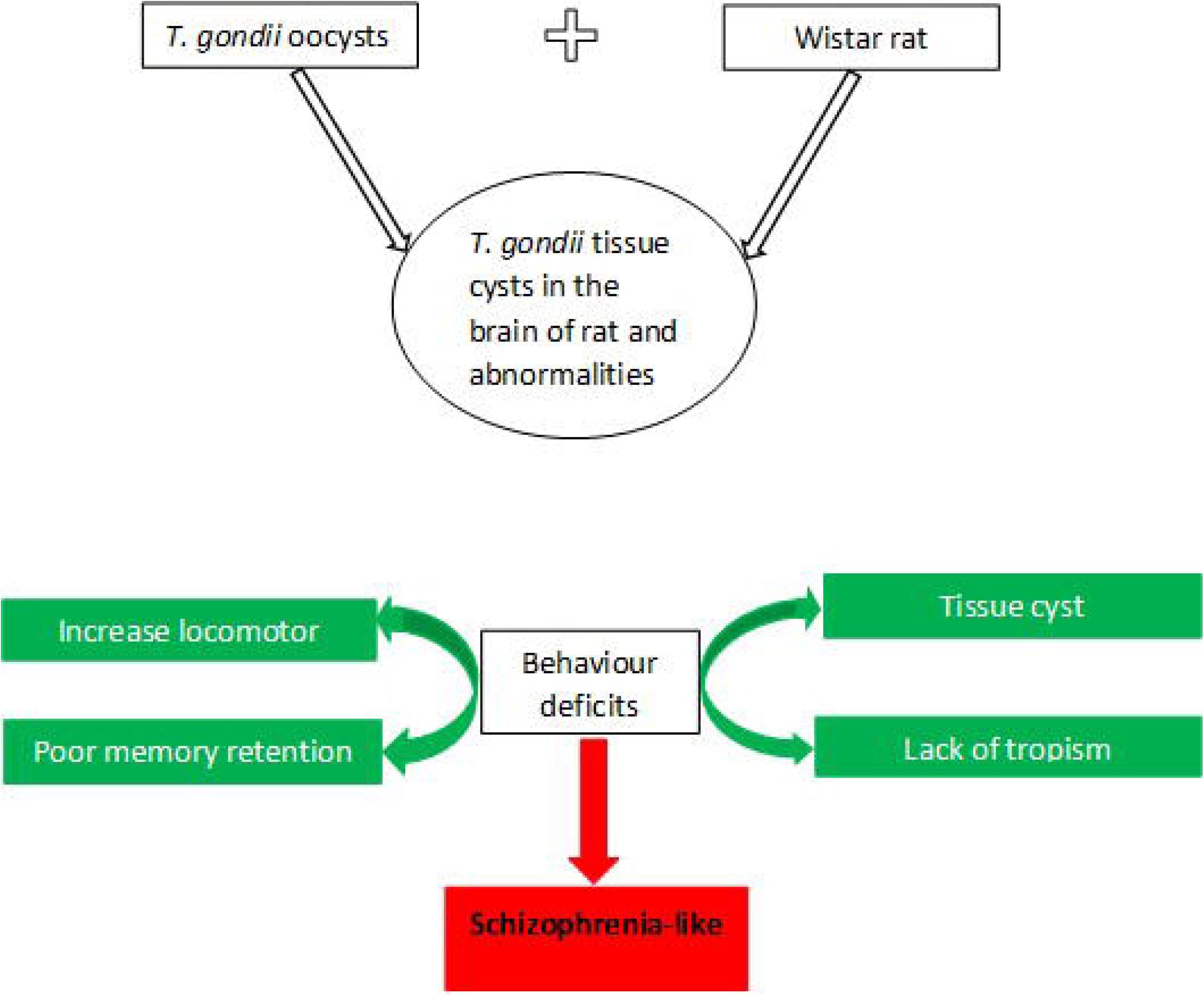

## HIGHLIGHTS

1. Malaysian strains of *T. gondii* from cat faeces are clonal with predominant type I.
2. There is an increase in locomotor activity in MK-801 and *T. gondii* infected groups compared to control.
3. Rats infected with *T. gondii* strains presented with poor memory retention, while learning remain intact.
4. There is wide distribution of *T. gondii* tissue cysts in the brain of rats infected with *T. gondii* without tropism.

